# Resolution of ribosomal stalling by ABCF ATPases YfmR and YkpA/YbiT

**DOI:** 10.1101/2024.01.25.577322

**Authors:** Hiraku Takada, Keigo Fujiwara, Gemma C. Atkinson, Chiba Shinobu, Vasili Hauryliuk

**Author notes:** to whom correspondence should be addressed: Huraku Takada, Vasili Hauryliuk.

## Abstract

Efficiency of protein synthesis on the ribosome is strongly affected by the amino acid composition of the assembled amino acid chain. Challenging sequences include proline-rich motifs as well as highly positively and negatively charged amino acid stretches. Members of the F subfamily of ABC ATPases (ABCFs) have been long hypothesised to promote translation of such problematic motifs. In this study we have applied genetics and reporter-based assays to characterise the four housekeeping ABCF ATPases of *Bacillus subtilis*: YdiF, YfmM, YfmR/Uup and YkpA/YbiT. We show that YfmR cooperates with the translation factor EF-P that promotes translation of Pro-rich motifs. Simultaneous loss of both YfmR and EF-P results in a dramatic growth defect. Surprisingly, this growth defect can be largely suppressed though overexpression of an EF-P variant lacking the otherwise crucial 5-amino-pentanolylated residue K32. Using *in vivo* reporter assays, we show that overexpression of YfmR can alleviate ribosomal stalling on Asp-Pro motifs. Finally, we demonstrate that YkpA/YbiT promotes translation of positively and negatively charged motifs but is inactive in resolving ribosomal stalls on proline-rich stretches. Collectively, our results provide insights into the function of ABCF translation factors in modulating protein synthesis in *B. subtilis*.

## INTRODUCTION

Protein synthesis on the ribosome is assisted by an array of dedicated protein factors that participate in all steps of translation: initiation, elongation, termination and recycling. The most well-studied group of ribosome-associated factors is translational GTPases (1–3). These factors promote the ‘core’ activities of the ribosome: bacterial initiation factor 2, IF2, promotes correct positioning of the initiator fMet-tRNA_i_, elongation factors EF-Tu and EF-G assist the delivery of aminoacyl-tRNA and catalyse ribosomal translocation, respectively, and, acting together with Ribosome Recycling Factor, RRF, EF-G splits the ribosome into subunits after the polypeptide is completed.

While translational GTPases all bind to the ribosomal A (acceptor) site, multiple ‘accessory’ factors act in the E (exit) site. Bacterial elongation factor P, EF-P, accesses the ribosomal peptidyl transferase center, PTC, to relieve ribosomal stalling on proline-rich motifs (4–8). PTC stimulation by EF-P requires posttranslational modification of the conserved lysine residue, with specific modifications differing in different bacterial lineages (9–13). In *Bacillus subtilis* EF-P is modified with a 5-aminopentanol moiety at Lys32 (11), via a multistep assembly pathway that relies on several enzymes: GsaB, YnbB, YmfI, YaaO, YfkA and YwlG (14). GsaB, YnbB and YmfI directly catalyse the EF-P modification while YaaO, YfkA and YwlG are believed to play an indirect role though supporting synthesis of the substrate (14). EF-P is not essential in *B. subtilis,* nor in *Escherichia coli* (14,15). EF-P loss results in a pleiotropic phenotype, which in *B. subtilis* involves compromised sporulation (due to the reduced expression of the Spo0A transcription factor) (16) and swarming mobility (due to the reduced expression of multiple swarming mobility-associated proteins, including FliP and FlhP) (11). However, EF-P is essential in other bacterial species such as *Neisseria meningitidis* (17), and the eukaryotic EF-P orthologue, eIF5A, is essential in yeast (18) and flies (19). Notably, in addition to its role in promoting translation elongation on proline-rich stretches, eIF5A also plays a crucial role in translation termination (20); no similar function has been shown for EF-P.

The F subfamily of ABC ATPases (ABCFs) comprises another group of E-site-binding translation factors in bacteria (21,22). The family includes both antibiotic resistance (ARE) factors as well as housekeeping proteins that assist protein synthesis and ribosome assembly (23–27). The *B. subtilis* genome encodes five ABCFs: a dedicated antibiotic resistance factor VmlR (28) and housekeeping factors YdiF, YfmM, YfmR/Uup and YkpA/YbiT (25). The exact functions of housekeeping ABCFs are unclear. However, multiple lines of evidence suggest that, similarly to ARE ABCF that resolve ribosome stalling caused by antibiotics (29–33), housekeeping ABCFs are expected to reach into the PTC with their P-site tRNA interaction motif (PtIM) domain in order to resolve other stalling events in an NTPase-dependent manner (23,24,34) (**Figure 1**). *E. coli* EttA is by far the best characterised housekeeping ABCF, with structural and biochemical evidence indicating a role in the regulation of the first rounds of translation elongation (23,24,34). The EttA subfamily evolved from the diversity of the Uup ABCF subfamily (25). Several studies strongly suggest a non-ribosomal role for Uup in resolving DNA repair intermediates (35,36) and transposon excision (37). However, overexpression of Uup suppresses the cold sensitivity and ribosome assembly defects of an *E. coli* strain lacking the accessory translational GTPase BipA (29), suggesting a direct role for Uup in translation or/and ribosome assembly. The ribosomal function of Uup is further supported by specific inhibition of protein synthesis upon expression of the ATPase-deficient (EQ_2_) variant due to non-productive association of the Uup-EQ_2_ variant with the ribosome (29,38). Ectopically overexpressed ABCF-EQ_2_ variants preferentially bind to the vacant E site of 70S initiation complexes (IC) (23,24) which has been successfully exploited for immunoaffinity-based purification of ABCF-EQ_2_:IC complexes for structural studies (32,33,39,40).

**Figure 1.**
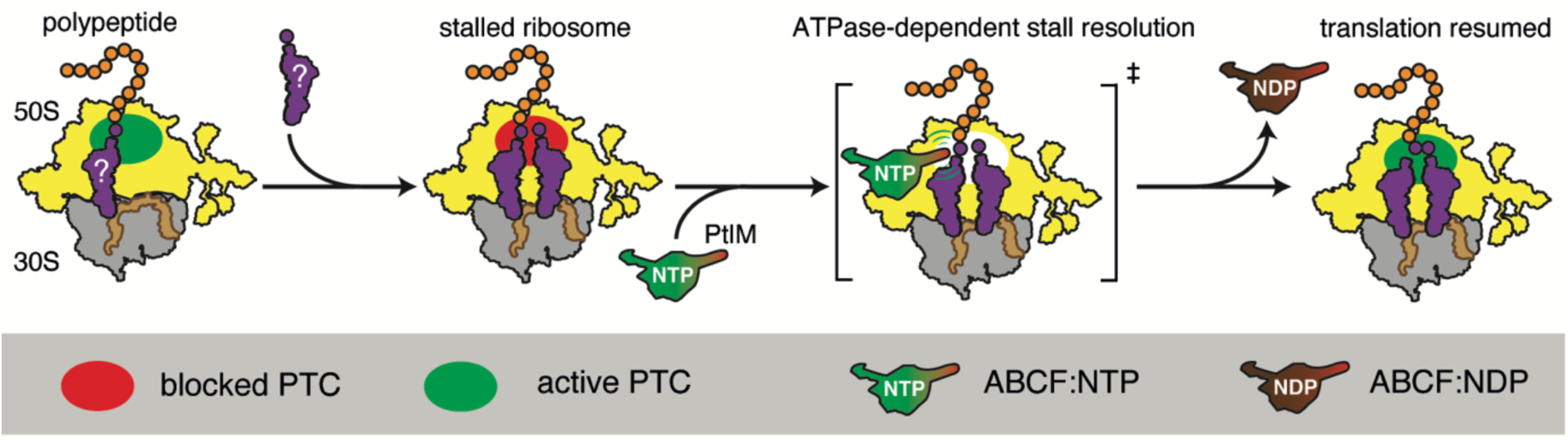
Generalised model for ribosomal rescue by housekeeping ABCF ATPases. Translation though ‘challenging’ amino acid motifs stalls the ribosome, inactivating the PTC. Housekeeping ABCF binds to the ribosomal E site and accesses the PTC with its P-site tRNA interaction motif (PtIM) domain. Following the NTPase-dependent reset of the PTC by the factor, the ABCF departs from the E site; translation resumes. The exact nature of the stalls resolved by the individual ABCFs is currently unclear.

We have characterised the potential functional overlap between the two classes of E-site-inspecting factors in *B. subtilis*: EF-P and housekeeping ABCFs. While the two classes cannot operate on the ribosome simultaneously due to a steric clash, we show genetic and functional interactions between EF-P and YfmR, with functional assays demonstrating that *B. subtilis* YfmR is able to resolve ribosome stalling on Asp-Pro motifs. Furthermore, we demonstrate that YkpA/YbiT promotes translation of EF-P-insensitive positively and negatively charged motifs.

## MATERIALS AND METHODS

### Phylogenetic analysis

Representative sequences were selected from Murina *et al.* (25). Sequences were aligned with MAFFT v6.861b with the L-INS-i strategy (41) and visualised in AliView (42). Positions with 50% gaps were removed with trimAl v. 1.4.rev6 (43). Maximum likelihood phylogenetic analysis was carried out with IQ-Tree v. 2.2.2.6 (44) at the IQ-Tree web server (45) with the best fit model selected by the program (Q.pfam+I+G4), SH-aLRT and ultrafast bootstrap and branch support testing (1000 replicates) (46).

### Construction of bacterial strains and plasmids

The strains, plasmids and oligonucleotides used in this study as well as description of strain construction are provided in **Supplementary Table 1**. Plasmids were constructed by standard cloning methods: PCR, PrimeSTAR mutagenesis (Takara), and Gibson assembly. Marker-less gene deletion mutants of *efp* (BCHT209), *ydiF* (BCHT212), *yfmM* (BCHT213), *yfmR* (BCHT214), *ykpA* (BCHT215), *gsaB* (BCHT332), *yaaO* (BCHT333), *yfkA* (BCHT334), *ymfI* (BCHT335) and *ynbB* (BCHT336) were constructed by excising the antibiotic resistance cassette by the Cre-loxP system as described previously (47). Briefly, *B. subtilis* strains were transformed with pMK2, a pLOSS*-based Ts plasmid harbouring *cre*. To select for the excision of the resistance marker flanked by *loxP*, the resulting strains were grown overnight at 37 °C on LB agar medium supplemented with 1 mM isopropyl β-D-1-thiogalactopyranoside (IPTG) and 100 μg/mL spectinomycin. Finally, to promote the loss of the pMK2 plasmid, the strains were grown overnight at 37 °C on LB agar medium without spectinomycin. The loss of pMK2 was confirmed by the absence of spectinomycin resistance.

### Sucrose gradient fractionation and immunoblotting

The experiments were performed as described previously (48). Briefly, *B. subtilis* strains were grown at 37 °C in 40 mL LB cultures until the OD_600_ of 0.8, cells collected by centrifugation and dissolved in 0.5 mL of HEPES:Polymix buffer [5 mM Mg(OAc)_2_] (48), lysed by FastPrep homogenizer (MP Biomedicals) and the resultant lysates clarified by centrifugation. 10 A_260_ units of each extract were loaded onto 10%-35% (w/v) sucrose density gradients prepared in HEPES:Polymix buffer [5 mM Mg(OAc)_2_] and the gradients were resolved by ultracentrifugation at 36 000 rpm for 3 h at 4 °C. Both preparation and fractionation of gradients was done using Biocomp Gradient Station (BioComp Instruments); A_260_ was used as a readout during the fractionation.

### Ribosome stalling reporter assay

Ribosome stalling reporters were based on the reporters developed by Chadani and colleagues (49). Reporters were expressed under the control of P*_hy-spank_* IPTG-inducible promoter (50) from a self-replicated pHT01-based plasmid carrying a kanamycin resistance marker (51). ABCF- and EF-P-coding genes were cloned on the pSHP2 plasmid (provided by Dr. Henrik Strahl von Schulten) under the control of P*_xy_* xylose-inducible promoter and integrated into the *amyE* locus. Individual reporter plasmids were amplified by EquiPhi29 polymerase (Thermo Fisher Scientific) and transformed into recipient *B. subtilis* strains. The resulting strains were grown overnight at 37 °C on LB plates supplemented with 3 µg/mL kanamycin. After isolating single colonies twice on LB plates supplemented with 3 µg/mL kanamycin, fresh colonies of *B. subtilis* harbouring reporter plasmids were used to inoculate 1-mL LB medium cultures dispensed into plastic 96 deep-well plates (Treff Lab). The cultures were grown at 30 °C for 18 h with shaking at 1200 rev per min using DWMax M·BR-034P constant temperature incubator shaker (Taitec). 20 µL of individual overnight cultures were then used to inoculate 1 mL cultures (LB supplemented with 3 µg/ml kanamycin as well as inducers: 1 mM IPTG and 0.3% xylose) dispensed into plastic 96 deep-well plates. 1-mL experimental cultures were grown at 37 °C with shaking until OD_600_ of 1.0, 0.75 mL aliquots collected, combined with 83 µL of 50% TCA and kept on ice for 5 min. After centrifugation at 13 500 rpm for 2 min at 4 °C, cell pellets were resuspended in 500 µL of 0.1 M Tris-HCl (pH 6.8). After one more round of 2-min centrifugation at 13 500 rpm at 4 °C, cell pellets were resuspended in 50 µL of lysis buffer (0.5 M sucrose, 20 mM MgCl_2_, 1 mg/ml lysozyme, 20 mM HEPES:NaOH, pH 7.5) and incubated at 37 °C for 10 min. Next, an equal volume of 2× SDS sample buffer (4% SDS, 30% glycerol, 250 mM Tris pH 6.8, 1 mM DTT, saturated bromophenol blue) was added, and the lysates were denatured at 85 °C for 5 min. Proteins were resolved on 11% SDS-PAGE and transferred to a PVDF membrane. GFP-tagged proteins were detected using anti-GFP (Wako, mFX75, 1:5 000 dilution) antibodies combined with Goat Anti-Mouse IgG (H+L) HRP Conjugate (Bio-Rad). Images were acquired using Amersham Imager 600 (GE Healthcare) luminoimager and analysed in ImageJ (52). The stalled fraction was quantified by dividing the stalled (short) product signal by the total signal (short and full-length combined). All experiments were performed as three biological replicates; quantification is shown as mean ± standard deviation.

## RESULTS

### B. subtilis YfmR is a member of the Uup/EttA ABCF clade

YfmR is classified with ABCF Hidden Markov models as a member of the Uup subfamily (25). The Uup subfamily is not monophyletic, but rather is paraphyletic to the EttA subfamily, which arose from an Uup-like progenitor (**Figure 2**, (25)). Uup subfamily members typically carry a C-terminal domain (CTD), which is absent in EttA, suggesting this domain was lost after the *uup* gene duplication that gave rise to EttA. The monophyly of the Uup+EttA clade is strongly supported (99.7% SH-aLRT and 100% bootstrap support).

**Figure 2.**
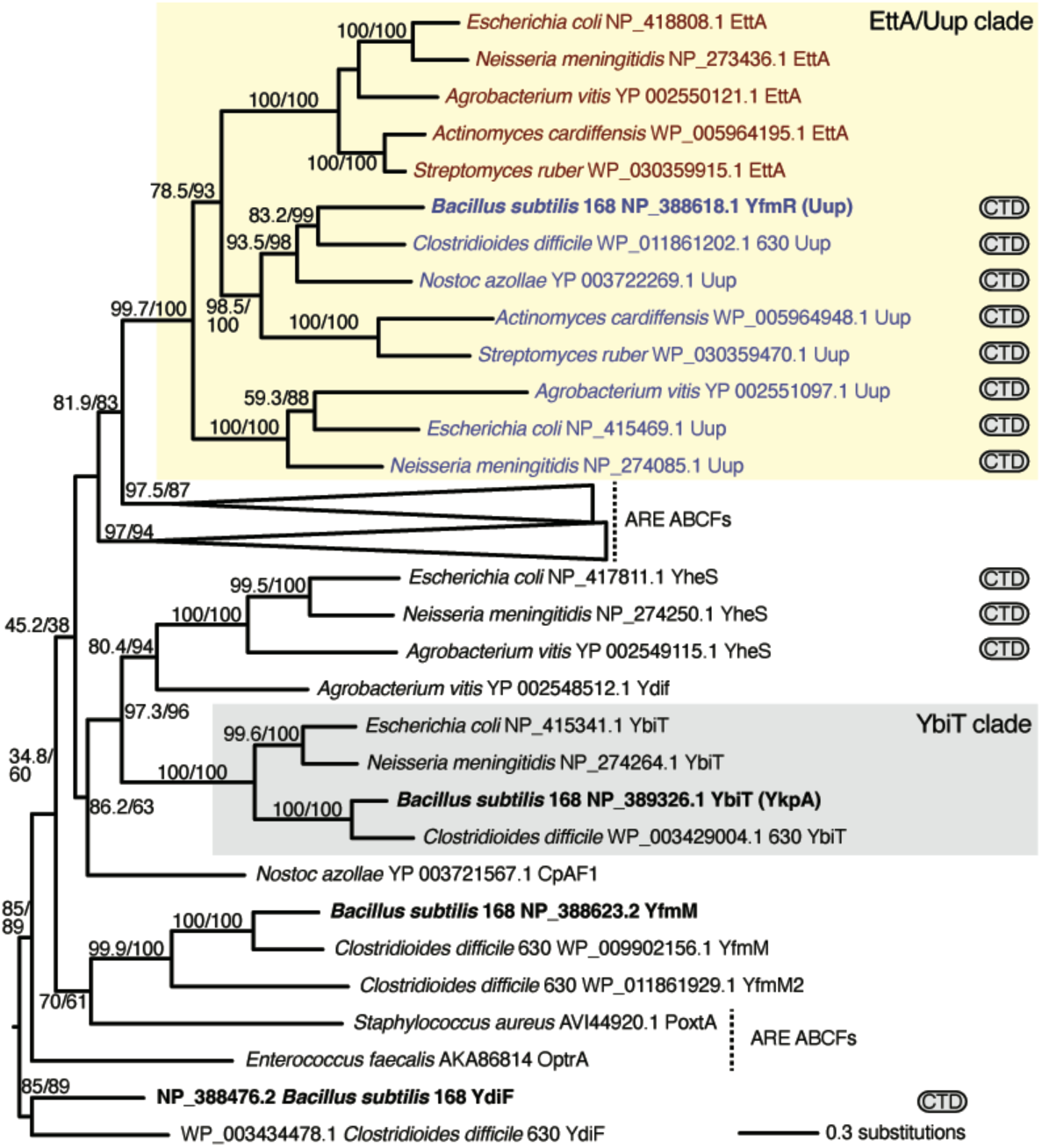
*B. subtilis* YfmR is an Uup ABCF subfamily member within the EttA/Uup clade. Maximum likelihood phylogeny of ABCF protein representatives. *B. subtilis* housekeeping ABCFs YfmR, YdiF, YfmM and YkpA (YbiT) are shown in bold. The CTD logo indicates the presence of a C-terminal domain. Numbers in parentheses are SH-aLRT support (%) / ultrafast bootstrap support (%). Only branches with more than 60% bootstrap support are labelled. Branch length is proportional to the number of substitutions as per the lower right key.

### Simultaneous disruption of yfmR and efp results in a synthetic growth defect

We took a genetic approach to probe the functional interactions between housekeeping ABCFs and EF-P in the *B. subtilis* 168 strain. While genomic disruptions of individual ABCF genes in the wild-type background do not affect *B. subtilis* growth on LB medium at the optimal temperature (37 °C), deletion of *yfmR* – but not any of the other three housekeeping ABCFs – results in severe growth defect in the Δ*efp* background (**Figure 3A**). Sucrose gradient centrifugation experiments reveal the low abundance of polysomes as well as accumulation of 40S ribosome assembly intermediates in the Δ*efp* Δ*yfmR* double deletion strain, consistent with perturbed protein synthesis (**Supplementary Figure S1**). Finally, no genetic interaction was observed for *efp* and poorly understood translational GTPases *bipA* and *lepA* (**Figure 3A**), suggesting, expectedly, that these factors are not operating together with EF-P.

**Figure 3.**
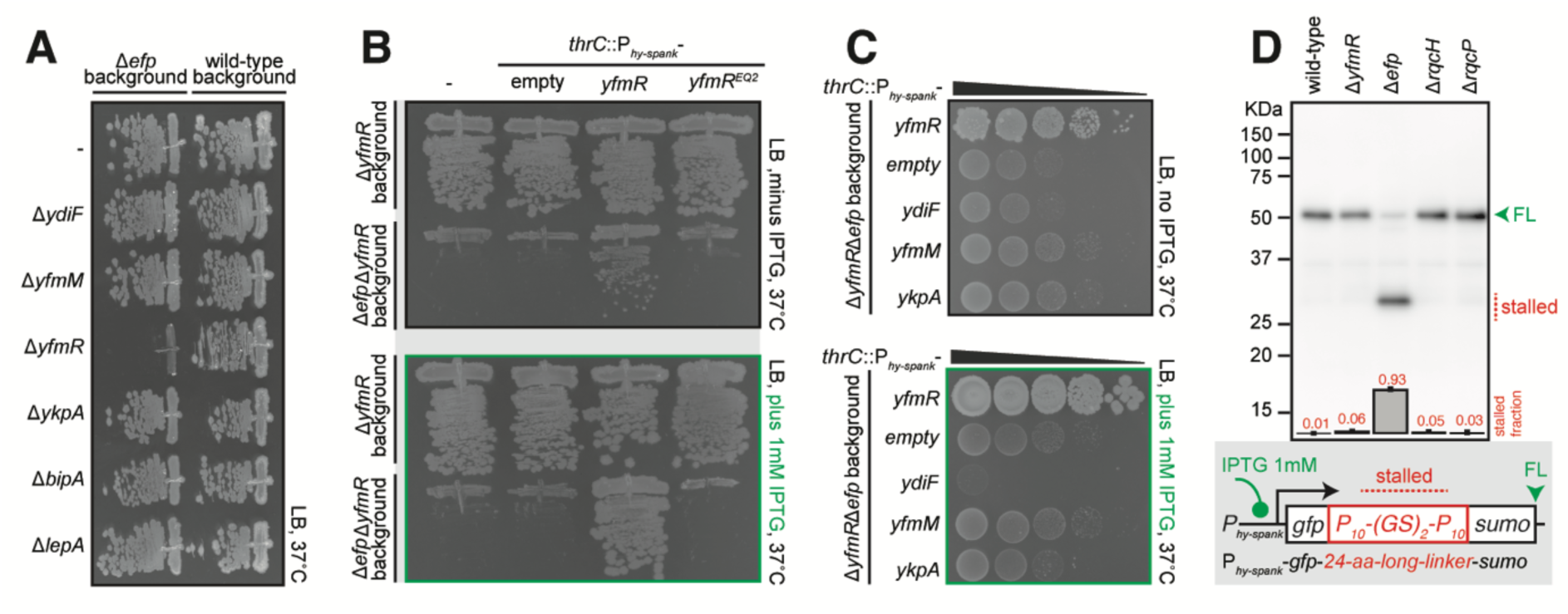
Simultaneous loss of *yfmR* and *efp* results in a dramatic growth defect. (A) Effects of mutations targeting housekeeping ABCFs and non-essential translational GTPases BipA and LepA on the growth of wild-type and Δ*efp B. subtilis*. The Δ*ydiF* (strains BCHT174 and BCHT195; wt and Δ*efp* background, respectively), Δ*yfmM* (strains BCHT171 and BCHT192), Δ*yfmR* (strains BCHT170 and BCHT191), Δ*ykpA* (strains BCHT170 and BCHT191), Δ*bipA* (strains BCHT172 and BCHT193) and Δ*lepA* (strains BCHT173 and BCHT194) *B. subtilis* were grown on solid LB medium at 37 °C. Wild-type *B. subtilis* and the isogenic Δ*efp* mutant (strain BCHT175) were streaked as controls. (B) Overexpression of the ATPase-deficient YfmR-EQ_2_ mutant does not rescue the growth defect of Δ*yfmR*Δ*efp B. subtilis*. *B. subtilis* strains expressing wild-type YfmR (BCHT385 and BCHT389; Δ*yfmR* or Δ*yfmR*Δ*efp* backgrounds, respectively) or YfmR-EQ_2_ (strains BCHT386 and BCHT390) were grown on solid LB medium with (*lower panel*) or without (*upper panel*) 1 mM IPTG. Δ*yfmR* mutant (strains BCHT181 and BCHT170) and Δ*efp*Δ*yfmR* mutant (strains BCHT187 and BCHT191), both with or without integration of the empty vector were streaked as controls. (C) Effects of housekeeping ABCF overexpression on the growth of Δ*efp*Δ*yfmR B. subtilis*. YfmR (strain BCHT385 and BCHT389), YdiF (strain BCHT440 and BCHT435), YkpA (strain BCHT441 and BCHT436) and YfmM (strain BCHT442 and BCHT437) were overexpressed in either Δ*yfmR* or Δ*yfmR*Δ*efp B. subtilis* growing on solid LB medium with (*lower panel*) or without (*upper panel*) 1 mM IPTG. (D) Polyproline stalling reporter (GFP-P_10_-(GS)_2_-P_10_-SUMO, pCHT55) detects ribosomal stalling in Δ*efp* but not Δ*yfmR B. subtilis*. The reporter was expressed in wild-type, Δ*yfmR* (strain BCHT214), Δ*efp* (strain BCHT214), Δ*rqcH* (strain BCHT58) or Δ*rqcP* (strain BCHT56) *B. subtilis* and detected with anti-GFP antibodies. The full-length product is indicated with a green arrowhead and the stalled product is indicated with a red dotted line. Fraction of the stalled (short) product was quantified from three independent biological replicates and shown as mean ± standard deviation.

We have complemented the double deletion Δ*efp* Δ*yfmR* strain with either wild-type or the ATPase-deficient EQ_2_ variant of YfmR expressed under the control of IPTG-inducible P*_hy-spank_* promoter (50). Even in the absence of IPTG, leaky expression of the wild-type protein partially suppressed the growth defect; addition of 1 mM IPTG resulted in full suppression (**Figure 3B**). Low-level expression of the ATPase-deficient YfmR-EQ_2_ driven by the native Shine-Dalgarno motif fails to complement, consistent with the ATPase activity being essential (**Figure 3B**). Next, we tested whether ectopic overexpression of housekeeping ABCFs could suppress the growth defect of the Δ*efp* Δ*yfmR* strain. While P*_hy-spank_*-driven overexpression of YfmM and YkpA/YbiT has no effect, overexpression of YdiF further exacerbates the growth defect of Δ*efp* Δ*yfmR B. subtilis* (**Figure 3C**). A genetic interaction between *efp* and *ydiF* has previously been shown earlier by Hummels and Kearn who have demonstrated that the swarming defect of the Δ*efp B. subtilis* strain can be suppressed by loss-of-function mutations in *ydiF* (53). Notably, in the absence of IPTG, low-level leaky expression of YfmM and YkpA partially suppresses the growth defect of the Δ*efp* Δ*yfmR* strain, which could suggest partial functional redundance between YfmR and these two ABCF ATPases.

### YfmR is not essential for efficient translation of polyproline stretches in *efp^+^ B. subtilis*

There can be several alternative explanations for the strong genetic interaction between *efp* and *yfmR*. One possibility is that EF-P and YfmR are both required for translation of proline-rich motifs, providing partially redundant solutions to this stalling problem. To test this hypothesis we used a homopolymeric polyproline stalling reporter based on that developed by Chadani and colleagues (49). The reporter gene encodes an N-terminal GFP and C-terminal SUMO tag linked by a 24-amino-acid-long linker with a sequence of P_10_-(GS)_2_-P_10_; two 10-amino-acid-long stalling motifs connected by a flexible ‘joint’ that is not stalling-prone. The gene encoding the reporter was cloned on a self-replicated plasmid pHT01-based plasmid (51) and the expression was driven by IPTG-inducible P*_hy-spank_* promoter. While the full-length version of the reporter construct was efficiently produced in the wild-type *B. subtilis*, only a fraction of the full-length product is synthesised in the Δ*efp* strain, with the majority of the ribosomes stalling and generating a short version of the reporter (**Figure 3D**). The Δ*yfmR* strain behaved like the wild-type, with no short versions of the reporter being produced. Therefore, we concluded that YfmR is unlikely to be involved in translation of strongly stalling polyproline stretches and that EF-P and YfmR synergise in other, yet-undefined stalling motifs.

### EF-P variants lacking the K32 residue or its 5-aminopentanol modification can still efficiently suppress of the growth defect of Δefp Δ*yfmR B. subtilis*

With the notable exception of Actinobacterial EF-P (54), the posttranslational modification of a conserved lysine residue is crucial for the factor’s functionality in resolving polyproline stalling, both in living cells and a reconstituted biochemical system (6,8,10,14). However, it is conceivable that the modification is not essential for the hypothetical activity on which EF-P and YfmR work together. To probe this hypothesis, we tested whether EF-P lacking the modification of conserved K32 residue – or the K32 residue altogether – can suppress the synthetic growth defect of Δ*efp* Δ*yfmR B. subtilis*. Surprisingly, a Δ*yfmR B. subtilis* strain expressing the K32A EF-P variant does not phenocopy the severe growth defect of the Δ*efp* Δ*yfmR* double knockout (**Figure 4A**). Furthermore, systematic disruptions of the genes involved in the 5-aminopentanol modification of the K32 residue – *gsaB*, *yaaO*, *ynbB*, *yfkA* and *ywlG* (14) – do not strongly genetically interact with the *yfmR* disruption: none of the double-knockout strains display a severe growth defect either; a minor growth defect is detectable in Δ*yaaO* Δ*yfmR* (**Figure 4B**).

**Figure 4.**
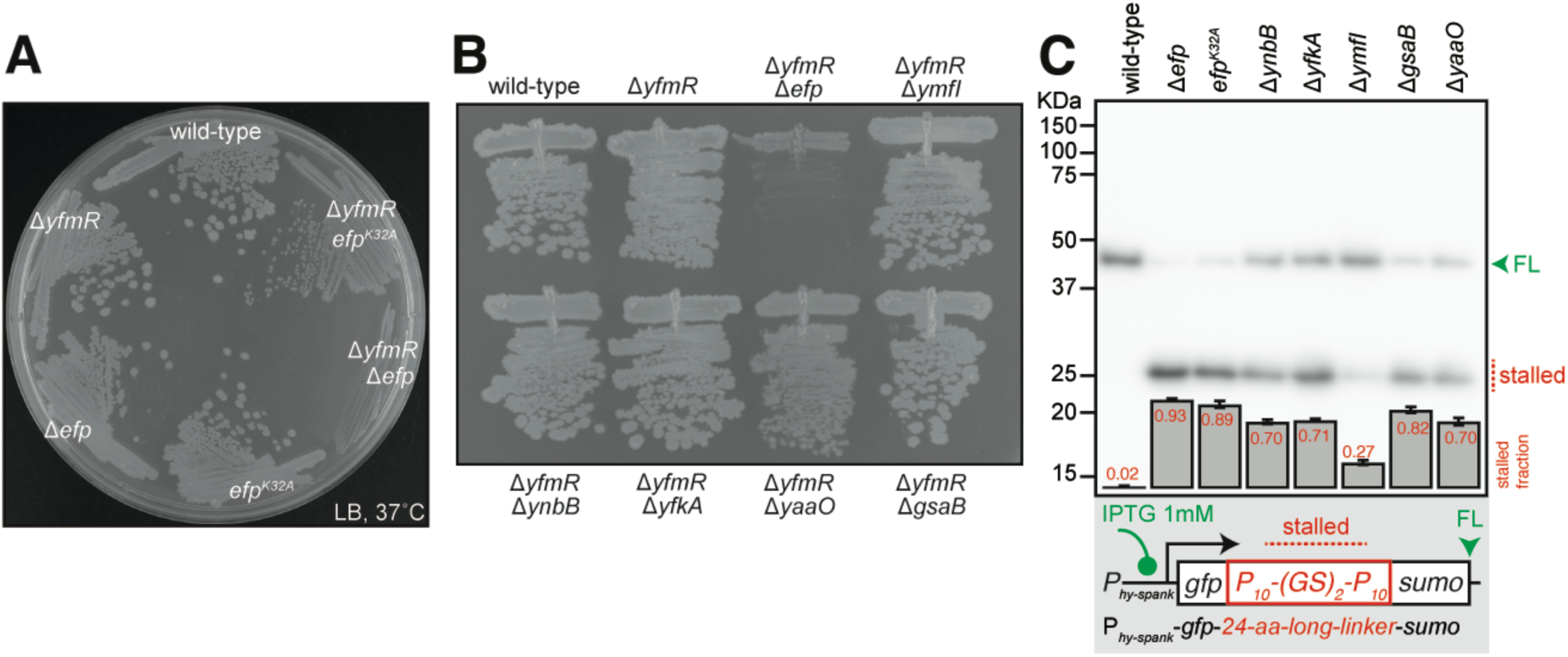
5-amino-pentanolylated residue K32 of EF-P is not essential for supporting a near-wild-type fitness of the Δ*yfmR B. subtilis*. (A) Effect of the *efp^K32A^* mutation on the negative genetic interaction between *yfmR* and *efp*. Wild-type, Δ*yfmR* (strain BCHT170), Δ*efp* (strain BCHT175), *efp^K32A^* (strain BCHT8765), Δ*yfmR efp^K32A^* (strain BCHT879) and Δ*yfmR*Δ*efp* (strain BCHT191) *B. subtilis* were grown on solid LB medium at 37 °C. (B) Effects of disruption of the genes involved in the 5-aminopentanol modification of the K32 residue of EF-P on the growth of Δ*yfmR B. subtilis*. Wild-type, Δ*yfmR* (strain BCHT170), Δ*yfmR*Δ*efp* (strain BCHT191), Δ*yfmR*Δ*ymfI* (strain BCHT396), Δ*yfmR*Δ*ynbB* (strain BCHT395), Δ*yfmR*Δ*yfkA* (strain BCHT394), Δ*yfmR*Δ*yaaO* (strain BCHT393) and Δ*yfmR*Δ*gsaB* (strain BCHT392) *B. subtilis* were grown on solid LB medium at 37 °C. (C) Effects of the K32A substitution and genetic disruption of the K32 5-aminopentanol modification on the ribosomal stalling on polyproline. GFP-P_10_-(GS)_2_-P_10_-SUMO reporter (pCHT55) was expressed in wild-type, Δ*efp* (strain BCHT214), *efp^K32A^* (strain BCHT765), Δ*ynbB* (strain BCHT336), Δ*yfkA* (strain BCHT334), Δ*ymfI* (strain BCHT335), Δ*gsaB* (strain BCHT332) or Δ*yaaO* (strain BCHT333) *B. subtilis* and detected with anti-GFP antibodies. The full-length product is indicated with a green arrowhead and the stalled product is indicated with a red dotted line. Fraction of the stalled (short) product was quantified from three independent biological replicates and shown as mean ± standard deviation.

We next used our polyproline stalling reporter [GFP-P_10_-(GS)_2_-P_10_-SUMO] to assess the effects of the K32A substitution – as well as disruption of the 5-aminopentanol modification pathway – on EF-P’s functionality in promoting translation elongation on polyproline stretches. The K32A substitution phenocopied the Δ*efp* strain (**Figure 4C**). This result is in good agreement with analogous *in vivo* assays by Rajkovic and colleagues who tested an array of different PPX stalling peptides such as PPW, PPG, PPP, PPR (11). Mutations in the enzymes implicated in the 5-aminopentanol modification of the K32 residue strongly – although not as completely as the K32A substitution – compromised EF-P’s activity (**Figure 4C**). The weakest effect was observed in the case of disruption of YfmI, an enzyme which catalyses the last step in EF-P modification, the reduction EF-P-5 aminopentanone to EF-P-5 aminopentanol (55).

### Overexpression of YfmR/Uup alleviates the ribosomal stalling on Asp-Pro motifs in Δ*efp B. subtilis*

We next tested whether overexpression of YfmR can improve the ability of the *B. subtilis* translational apparatus to synthesise challenging motifs. Inspired by the work by Chadani and colleagues (49), we used a series of diverse GFP-*linker*-SUMO reporters with different linker sequences. Specifically, we tested homopolymeric P_10_- (GS)_2_-P_10_ and D_10_-(GS)_2_-D_10_ as well as ‘mixed’ (DP)_5_-(GS)_2_-(DP)_2_. The two proline-rich linkers are expected to specifically cause ribosomal stalling in Δ*efp B. subtilis* while the highly negatively charged poly-Asp motif is in general challenging for translation (49). Finally, the A_10_-(GS)_2_-A_10_ motif was used as a negative control as no stalling is expected in wild-type and Δ*efp B. subtilis*. The FLAG-tagged versions of either wild-type or functionally compromised proteins (ATPase-deficient EQ_2_ variants) were expressed under the control of a xylose-inducible P*_xy_* promotor. In this case YfmR was expressed under the control of strong Shine-Dalgarno motif, and this strong expression of YfmR-EQ_2_ is associated with a growth defect, most likely due to non-productive ribosomal association inhibiting translation. We have observed analogous effects in the case of ATPase-deficient housekeeping ABCFs in *E. coli* (25). Overexpression of either wild-type or K32A-substituted EF-P was used as two additional controls, and the reporter assays were performed either in wild-type or a Δ*efp* genetic background.

As expected, no truncated versions of the control GFP-A_10_-(GS)_2_-A_10_-SUMO reporter are detectable regardless of the strain background and the protein expressed (**Figure 5A**). Near 100%-efficient stalling on P_10_-(GS)_2_-P_10_ in Δ*efp B. subtilis* is fully resolved upon overexpression of wild-type EF-P; overexpression of the K32A-substituted variant failed to resolve the stall (**Figure 5B**). While overexpression of wild-type YfmR did not restore the production of the full-length reporter, it resulted in formation of a longer stalled product (marked with a red asterisk on **Figure 5B**), indicative of a possible modest stimulatory effect. Expression of YfmR-EQ_2_ decreased both the full-length and stalled reporter signal, most likely due to translation inhibition caused by the factor being locked in the ribosomal E-site. Experiments with a weaker EF-P-sensitive staller, (DP)_5_-(GS)_2_-(DP)_2_, indicated the ability of YfmR to resolve ribosomal stalls on proline-rich motifs: expression of either EF-P or YfmR abrogated the stalled signal, and the effect was specific for wild-type factors (**Figure 5C**, **Supplementary Figure S2**). Finally, we tested whether overexpression of either EF- P or YfmR could overcome ribosomal stalling on acidic poly-Asp motifs (**Figure 5D**). In agreement with EF-P not being able to resolve the poly-Asp stalling, the strength of stalled signal was similar in wild-type and Δ*efp B. subtilis*. Overexpression of neither of the factors could resolve ribosomal stalling on the D_10_-(GS)_2_-D_10_ motif.

**Figure 5.**
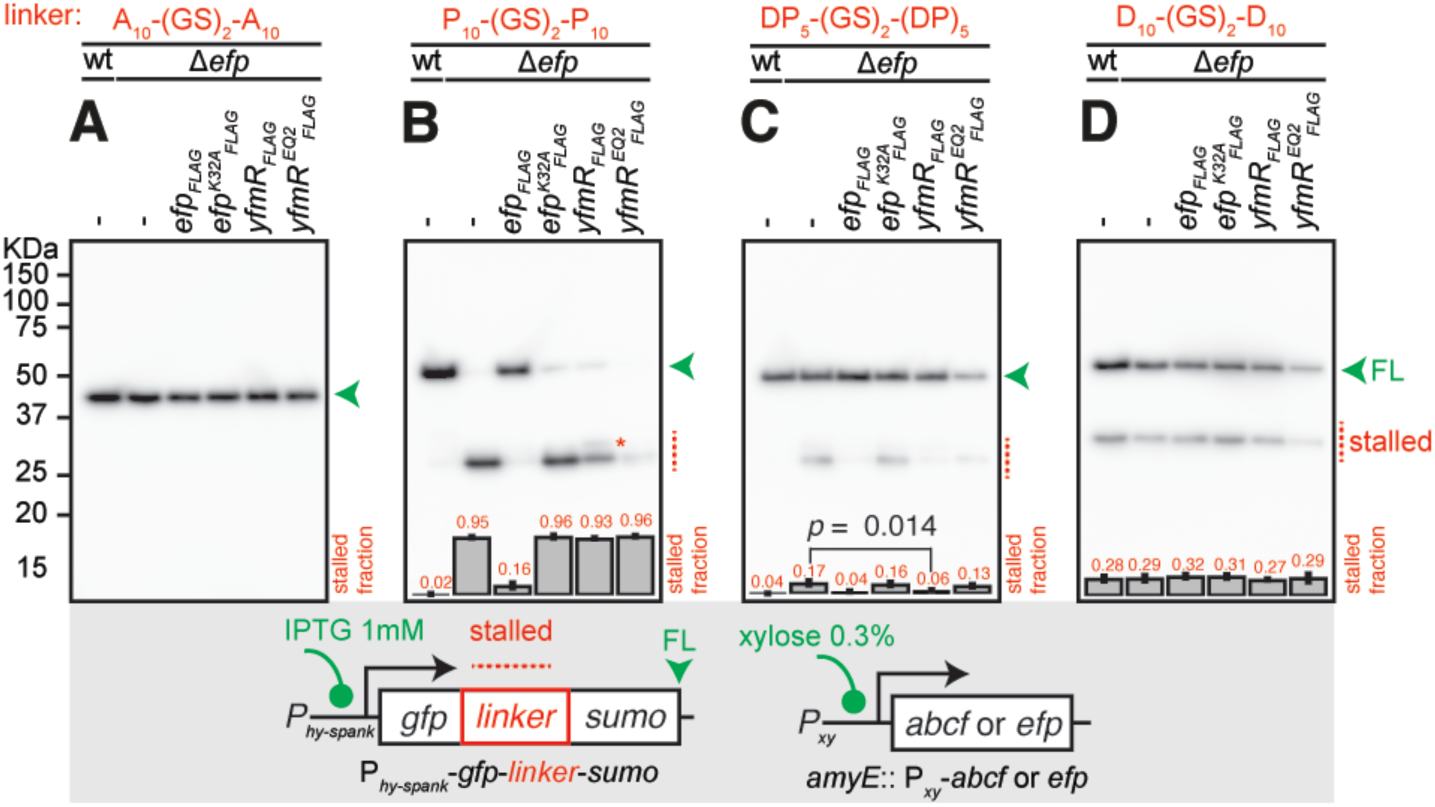
Overexpression of YfmR/Uup ABCF alleviates the ribosomal stalling on Asp-Pro motifs. Effects of EF-P and YfmR overexpression on ribosomal stalling on polyproline, polyaspartic acid as well as mixed Asp-Pro stalling motifs. GFP-A_10_-(GS)_2_-A_10_-SUMO (pCHT54) (**A**), GFP-P_10_-(GS)_2_-P_10_-SUMO (pCHT55) (**B**), GFP-(DP)_5_-(GS)_2_-(DP)_5_- SUMO (pCHT12) (**C**) and GFP-D_10_-(GS)_2_-D_10_-SUMO (pCHT15) (**D**) reporters were expressed in wild-type, Δ*efp* (strain BCHT214) as well as in Δ*efp B. subtilis* expressing either *efp_FLAG_* (strain BCHT1367), *efp^K32A^_FLAG_* (strain BCHT1368), *yfmR_FLAG_* (strain BCHT1369) or *yfmR^EQ2^_FLAG_* (strain BCHT1370) under the control of xylose promoter. All reporters were detected with anti-GFP antibodies. The full-length product is indicated with a green arrowhead and the stalled product is indicated with a red dotted line. A red asterisk indicates a larger stalled reporter product observed upon overexpression of YfmR-FLAG. Fraction of the stalled (short) product was quantified from three independent biological replicates and shown as mean ± standard deviation. An unpaired one-tailed Student’s t-test was used to compare Δ*efp* and Δ*efp* + *efp_FLAG_* groups on the panel (C). The effect size, measured as the ratio of sample means is 3.14 and the p-value is 0.014. The three individual experimental replicates of the panel (C) are shown on the **Supplementary Figure S2.**

B. subtilis *YkpA/YbiT promotes translation of positively and negatively charged motifs* Prompted by our results with YfmR, we decided to examine the possible involvement of all of the four *B. subtilis* ABCFs – YfmR, YdiF, YfmM and YkpA/YbiT – in translating diverse challenging sequences. As stretches of both highly positively and negatively charged amino acids can be challenging for the ribosome (49,56–59), we have also included polybasic [K_10_-(GS)_2_-K_10_ and R_10_-(GS)_2_-R_10_] motifs as well as an additional polyacidic [D_10_-(GS)_2_-D_10_] linker, respectively.

We tested all of the reporters listed above in wild-type and Δ*efp B. subtilis* as well as the four Δ*abcf B. subtilis* strains: Δ*yfmR,* Δ*ydiF,* Δ*ykpA* or Δ*yfmM.* As expected, all of the strains produce exclusively the full-length version of the GFP-A_10_-(GS)_2_-A_10_-SUMO reporter (**Figure 6A**). None of the ABCFs are crucial for translation of polyproline stretches in *efp*^+^ *B. subtilis*: while the short, stalled version of the GFP-P_10_- (GS)_2_-P_10_-SUMO reporter is dominant in Δ*efp B. subtilis*, only the full-length signal is detectable in all of the Δ*abcf* strains (**Figure 6B**). An analogous result was obtained with GFP-(DP)_5_-(GS)_2_-(DP)_2_-SUMO, although the strength of stalling in the Δ*efp* background is considerably weaker: the stalled form constitutes about 20% of the total signal (**Figure 6C**). Experiments with polybasic stallers yielded non-trivial results. In the case of K_10_-(GS)_2_-K_10_ linker we detected specific (but relatively weak) stalling in the Δ*ykpA* (Δ*ybiT*) strain (**Figure 6D**). While the R_10_-(GS)_2_-R_10_ motif was challenging for all of the tested strains, the strongest stalling signal was, again, observed in the case of Δ*ykpA B. subtilis*. Furthermore, weak Δ*ykpA*-specific stalling was observed in the case of E_10_-(GS)_2_-E_10_ polyacidic linker (**Figure 6F**). The polyacidic D_10_-(GS)_2_-D_10_ reporter was equally challenging for all of the tested strains (**Figure 6G**). Collectively, our results suggest that YbiT could be assisting the ribosome in negotiating charged amino acid patches.

**Figure 6.**
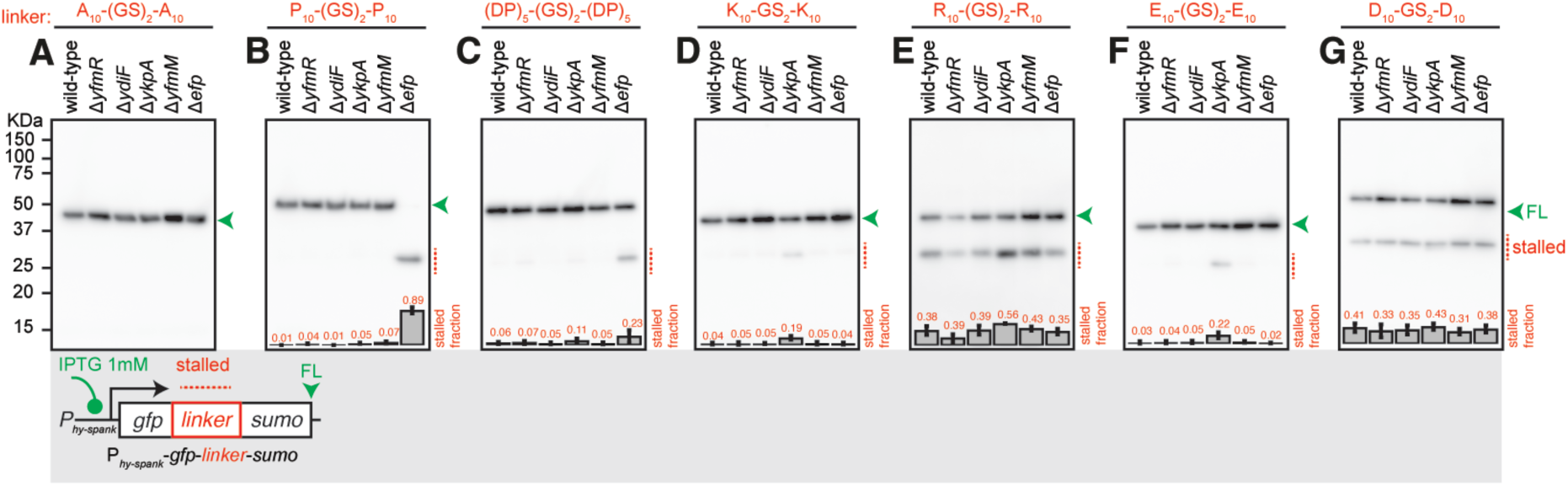
YkpA/YbiT loss results in mild ribosomal stalling on positively charged polylysine and negatively charged polyglutamic acid motifs. Effects of ABCF and EF-P gene disruption on ribosomal stalling on polyproline, Asp- Pro as well as negatively and positively charged homopolymeric motifs. GFP-A_10_- (GS)_2_-A_10_-SUMO (pCHT54) (**A**), GFP-P_10_-(GS)_2_-P_10_-SUMO (pCHT55) (**B**), GFP- (DP)_5_-(GS)_2_-(DP)_5_-SUMO (pCHT12) (**C**), GFP-K_10_-(GS)_2_-K_10_-SUMO (pCHT13) (**D**), GFP-R_10_-(GS)_2_-R_10_-SUMO (pCHT56) (**E**), GFP-E_10_-(GS)_2_-E_10_-SUMO (pCHT11) (**F**) and GFP-D_10_-(GS)_2_-D_10_-SUMO (pCHT15) (**G**) reporters were expressed in wild-type, Δ*yfmR* (strain BCHT214), Δ*ydiF* (strain BCHT212), Δ*ykpA* (strain BCHT215) and Δ*yfmM* (strain BCHT213). The full-length product is indicated with a green arrowhead and the stalled product is indicated with a red dotted line. All reporters were detected with anti-GFP antibodies. Fraction of the stalled (short) product was quantified from three independent biological replicates and shown as mean ± standard deviation.

## DISCUSSION

The exact molecular functions of housekeeping ABCFs have been ‘a riddle wrapped in a mystery inside an enigma’ of bacterial protein synthesis for a decade. Housekeeping ABCFs are expected to resolve ribosomal stalls – but what kind of stalls? Several recent *bioRxiv* preprints have provided important clues regarding the possible biological functions of both *E. coli* (38,60–62) and *B. subtilis* (63) factors. A study by Hong and colleagues has demonstrated that dCas9 knock-down of *yfmR* in Δ*efp B. subtilis* results a synthetic growth defect, increased ribosomal stalling on a pentaproline motif as well as accumulation of free 50S subunits. Furthermore, the authors showed that *E. coli* Uup can functionally replace YfmR/Uup in *B. subtilis*. An elegant biochemical study by Chadani and colleagues has demonstrated that in a reconstituted PURE*flex* protein synthesis system i) *E. coli* YbiT and EttA can suppress the premature termination on negatively charged polyacidic motifs and ii) *E. coli* Uup can alleviate ribosome stalling on polyproline stretches (61). Finally, in good agreement with Chadani *et al.*, Ousalem and colleagues have revealed the role of *E. coli* YbiT in alleviation of the ribosomal stalling acidic residues (62). All of these insights are well-aligned with our *in vivo* results with *B. subtilis*.

Despite the recent progress, our understanding of bacterial housekeeping ABCFs is still exceedingly incomplete. The contrast between, on one hand, the exceedingly strong and specific genetic interaction between *efp* and *yfmR* and, on the other hand, rather modest effects in stalling reporter assays is stark. Even more intriguing is the ability of the EF-P variant lacking the 5-amino-pentanolylated residue K32 to suppress the growth defect of the *efp* Δ*yfmR* strain. While the K32A EF-P variant is inefficient in resolving ribosomal stalling on polyproline, it is clearly competent in assisting YfmR in its biological function. It is possible that the two factors act (sequentially? simultaneous ribosomal association of the two E-site-binders is impossible) on as-yet-unidentified stalling motifs, and, unlike the ‘classical’ proline-rich motifs, EF-P is competent in resolving these without the need of the K32 residue. In the absence a ‘smoking gun’, our highly reductionist reporter approach is incapable of identifying such motifs. Therefore, it is essential to apply global approaches such as ribosome profiling (64) or 5PSeq (65) to uncover the physiologically-relevant targets of *B. subtilis* housekeeping ABCF ATPases. Once the native targets of YfmR and YkpA/YbiT are established, structure-functional studies will reveal the molecular mechanism of stall resolution by the ABCFs. Capitalising on the molecular insights into the mechanism of EF-P-mediated stimulation of PTC activity, modulation of EF-P concentration has been adapted as a strategy for improved efficiency of incorporation of non-canonical amino acids (66,67). It is possible that housekeeping ABCF ATPases could be useful for similar protein engineering applications in the future.

## ACKNOWLEDGMENTS

We thank Marcus J.O. Johansson and Daniel N. Wilson for valuable comments on the manuscript, Artyom A. Egorov for help with data analysis, Yuhei Chadani for helpful discussions as well as Machiko Murata and Naoko Muraki for technical support.

This work was supported by the Knut and Alice Wallenberg Foundation (project grant 2020-0037 to G.C.A. and V.H.), the Swedish Research Council (Vetenskapsrådet) grants (2019-01085 and 2022-01603 to G.C.A., 2021-01146 to V.H.), Crafoord foundation (project grant Nr 20220562 to V.H.), the Estonian Research Council (PRG335 to V.H.), Cancerfonden (20 0872 Pj to V.H.), postdoctoral grant from the Umeå Centre for Microbial Research, UCMR (to H.T.), JST, ACT X, Japan (JP1159335 to H.T.), MEXT, JSPS Grant-in-Aid for Scientific Research (grants 20H05926 and 21K06053 to S.C., 23K05017 to H.T., 21K15020 to K.F.), Institute for Fermentation, Osaka (grant G-2021-2-063 to S.C.).

---

**Supplementary Figure S1.**
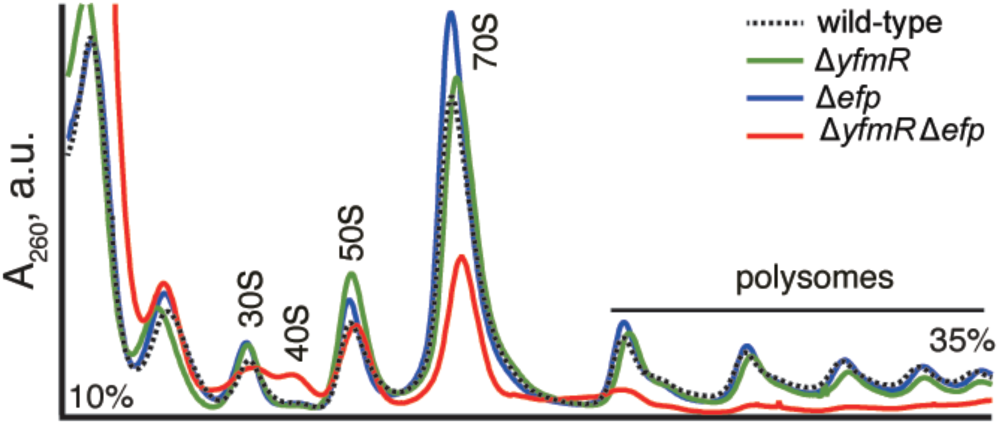
Sucrose gradient ultracentrifugation profiles of wild-type, Δ*yfmR,* Δ*efp as well as* Δ*efp* Δ*yfmR B. subtilis*. Sucrose gradient polysome analysis of wild-type, Δ*yfmR* (strain BCHT170), Δ*efp* (strain BCHT175) and Δ*efp*Δ*yfmR* (strain BCHT191) *B. subtilis*.

**Supplementary Figure S2.**
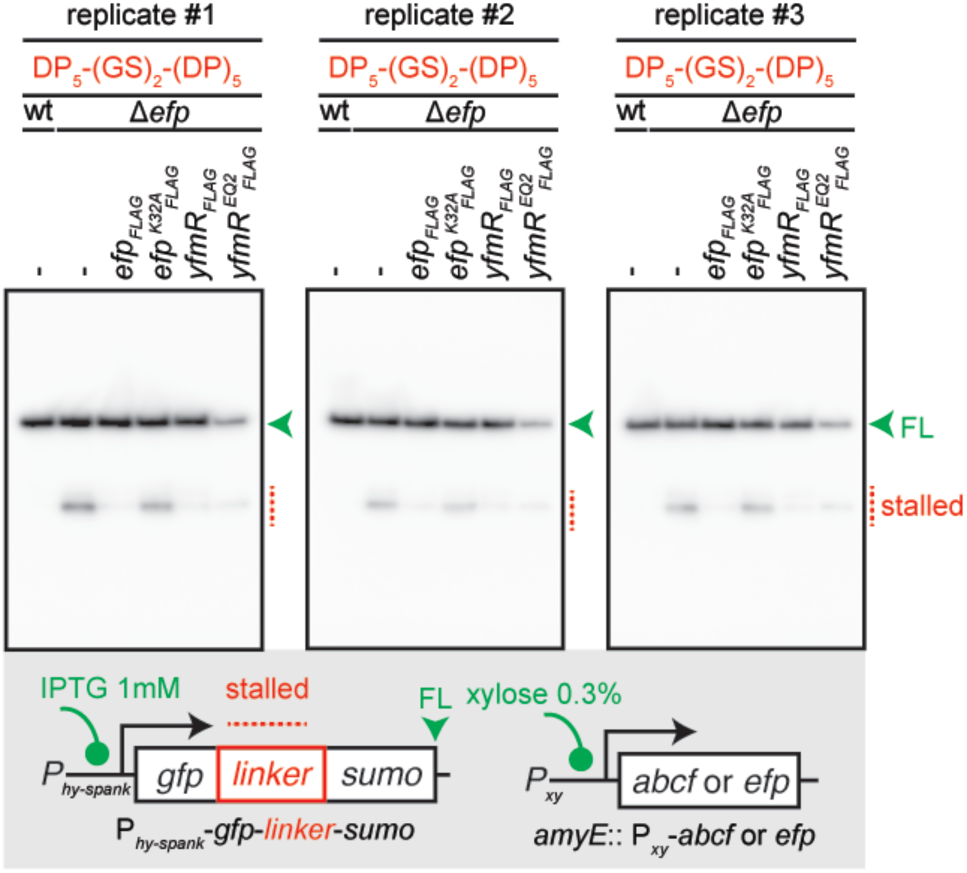
Overexpression of YfmR/Uup ABCF alleviates the ribosomal stalling on Asp-Pro motifs: tree individual biological replicates. Effects of EF-P and YfmR overexpression on ribosomal stalling on mixed Asp-Pro stalling motifs. The GFP-D_10_-(GS)_2_-D_10_-SUMO (pCHT15) reporter was expressed in wild-type, Δ*efp* (strain BCHT214) as well as in Δ*efp B. subtilis* expressing either *efp_FLAG_* (strain BCHT1367), *efp^K32A^_FLAG_* (strain BCHT1368), *yfmR_FLAG_* (strain BCHT1369) or *yfmR^EQ2^_FLAG_* (strain BCHT1370) under the control of xylose promoter and detected with anti-GFP antibodies. The full-length product is indicated with a green arrowhead and the stalled product is indicated with a red dotted line.

## Notes

### Competing Interest Statement

The authors have declared no competing interest.

## REFERENCES

1. Atkinson, G.C. (2015) The evolutionary and functional diversity of classical and lesser-known cytoplasmic and organellar translational GTPases across the tree of life. BMC Genomics, 16, 78.

2. Xu, B., Liu, L. and Song, G. (2021) Functions and Regulation of Translation Elongation Factors. Front Mol Biosci, 8, 816398.

3. Maracci, C. and Rodnina, M.V. (2016) Review: Translational GTPases. Biopolymers, 105, 463–475.

4. Huter, P., Arenz, S., Bock, L.V., Graf, M., Frister, J.O., Heuer, A., Peil, L., Starosta, A.L., Wohlgemuth, I., Peske, F. et al. (2017) Structural Basis for Polyproline-Mediated Ribosome Stalling and Rescue by the Translation Elongation Factor EF-P. Mol Cell, 68, 515–527 e516.

5. Doerfel, L.K., Wohlgemuth, I., Kothe, C., Peske, F., Urlaub, H. and Rodnina, M.V. (2013) EF-P is essential for rapid synthesis of proteins containing consecutive proline residues. Science, 339, 85–88.

6. Ude, S., Lassak, J., Starosta, A.L., Kraxenberger, T., Wilson, D.N. and Jung, K. (2013) Translation elongation factor EF-P alleviates ribosome stalling at polyproline stretches. Science, 339, 82–85.

7. Rajkovic, A. and Ibba, M. (2017) Elongation Factor P and the Control of Translation Elongation. Annu Rev Microbiol, 71, 117–131.

8. Peil, L., Starosta, A.L., Lassak, J., Atkinson, G.C., Virumae, K., Spitzer, M., Tenson, T., Jung, K., Remme, J. and Wilson, D.N. (2013) Distinct XPPX sequence motifs induce ribosome stalling, which is rescued by the translation elongation factor EF-P. Proc Natl Acad Sci U S A, 110, 15265–15270.

9. Peil, L., Starosta, A.L., Virumae, K., Atkinson, G.C., Tenson, T., Remme, J. and Wilson, D.N. (2012) Lys34 of translation elongation factor EF-P is hydroxylated by YfcM. Nat Chem Biol, 8, 695–697.

10. Lassak, J., Keilhauer, E.C., Furst, M., Wuichet, K., Godeke, J., Starosta, A.L., Chen, J.M., Sogaard-Andersen, L., Rohr, J., Wilson, D.N. et al. (2015) Arginine-rhamnosylation as new strategy to activate translation elongation factor P. Nat Chem Biol, 11, 266–270.

11. Rajkovic, A., Hummels, K.R., Witzky, A., Erickson, S., Gafken, P.R., Whitelegge, J.P., Faull, K.F., Kearns, D.B. and Ibba, M. (2016) Translation Control of Swarming Proficiency in *Bacillus subtilis* by 5-Amino-pentanolylated Elongation Factor P. J Biol Chem, 291, 10976–10985.

12. Park, J.H., Johansson, H.E., Aoki, H., Huang, B.X., Kim, H.Y., Ganoza, M.C. and Park, M.H. (2012) Post-translational modification by beta-lysylation is required for activity of *Escherichia coli* elongation factor P (EF-P). J Biol Chem, 287, 2579–2590.

13. Yanagisawa, T., Sumida, T., Ishii, R., Takemoto, C. and Yokoyama, S. (2010) A paralog of lysyl-tRNA synthetase aminoacylates a conserved lysine residue in translation elongation factor P. Nat Struct Mol Biol, 17, 1136–1143.

14. Witzky, A., Hummels, K.R., Tollerson, R., 2nd, Rajkovic, A., Jones, L.A., Kearns, D.B. and Ibba, M. (2018) EF-P Posttranslational Modification Has Variable Impact on Polyproline Translation in *Bacillus subtilis*. mBio, 9.

15. Tollerson, R., 2nd, Witzky, A. and Ibba, M. (2018) Elongation factor P is required to maintain proteome homeostasis at high growth rate. Proc Natl Acad Sci U S A, 115, 11072–11077.

16. Feaga, H.A., Hong, H.R., Prince, C.R., Rankin, A., Buskirk, A.R. and Dworkin,J. (2023) Elongation Factor P Is Important for Sporulation Initiation. J Bacteriol, 205, e0037022.

17. Yanagisawa, T., Takahashi, H., Suzuki, T., Masuda, A., Dohmae, N. and Yokoyama, S. (2016) *Neisseria meningitidis* Translation Elongation Factor P and Its Active-Site Arginine Residue Are Essential for Cell Viability. PLoS One, 11, e0147907.

18. Schnier, J., Schwelberger, H.G., Smit-McBride, Z., Kang, H.A. and Hershey, J.W. (1991) Translation initiation factor 5A and its hypusine modification are essential for cell viability in the yeast *Saccharomyces cerevisiae*. Mol Cell Biol, 11, 3105–3114.

19. Patel, P.H., Costa-Mattioli, M., Schulze, K.L. and Bellen, H.J. (2009) The Drosophila deoxyhypusine hydroxylase homologue nero and its target eIF5A are required for cell growth and the regulation of autophagy. J Cell Biol, 185, 1181–1194.

20. Schuller, A.P., Wu, C.C., Dever, T.E., Buskirk, A.R. and Green, R. (2017) eIF5A Functions Globally in Translation Elongation and Termination. Mol Cell, 66, 194–205 e195.

21. Gerovac, M. and Tampe, R. (2019) Control of mRNA Translation by Versatile ATP-Driven Machines. Trends Biochem Sci, 44, 167–180.

22. Ero, R., Kumar, V., Su, W. and Gao, Y.G. (2019) Ribosome protection by ABC-F proteins-Molecular mechanism and potential drug design. Protein Sci, 28, 684–693.

23. Böel, G., Smith, P.C., Ning, W., Englander, M.T., Chen, B., Hashem, Y., Testa, A.J., Fischer, J.J., Wieden, H.J., Frank, J. et al. (2014) The ABC-F protein EttA gates ribosome entry into the translation elongation cycle. Nat Struct Mol Biol, 21, 143–151.

24. Chen, B., Böel, G., Hashem, Y., Ning, W., Fei, J., Wang, C., Gonzalez, R.L., Jr., Hunt, J.F. and Frank, J. (2014) EttA regulates translation by binding the ribosomal E site and restricting ribosome-tRNA dynamics. Nat Struct Mol Biol, 21, 152–159.

25. Murina, V., Kasari, M., Takada, H., Hinnu, M., Saha, C.K., Grimshaw, J.W., Seki, T., Reith, M., Putrins, M., Tenson, T. et al. (2019) ABCF ATPases Involved in Protein Synthesis, Ribosome Assembly and Antibiotic Resistance: Structural and Functional Diversification across the Tree of Life. J Mol Biol, 431, 3568–3590.

26. Sharkey, L.K., Edwards, T.A. and O’Neill, A.J. (2016) ABC-F Proteins Mediate Antibiotic Resistance through Ribosomal Protection. MBio, 7, e01975.

27. Koberska, M., Vesela, L., Vimberg, V., Lenart, J., Vesela, J., Kamenik, Z., Janata, J. and Balikova Novotna, G. (2021) Beyond Self-Resistance: ABCF ATPase LmrC Is a Signal-Transducing Component of an Antibiotic-Driven Signaling Cascade Accelerating the Onset of Lincomycin Biosynthesis. mBio, 12, e0173121.

28. Ohki, R., Tateno, K., Takizawa, T., Aiso, T. and Murata, M. (2005) Transcriptional termination control of a novel ABC transporter gene involved in antibiotic resistance in *Bacillus subtilis*. J Bacteriol, 187, 5946–5954.

29. Murina, V., Kasari, M., Hauryliuk, V. and Atkinson, G.C. (2018) Antibiotic resistance ABCF proteins reset the peptidyl transferase centre of the ribosome to counter translational arrest. Nucleic Acids Res, 46, 3753–3763.

30. Crowe-McAuliffe, C., Graf, M., Huter, P., Takada, H., Abdelshahid, M., Novacek, J., Murina, V., Atkinson, G.C., Hauryliuk, V. and Wilson, D.N. (2018) Structural basis for antibiotic resistance mediated by the *Bacillus subtilis* ABCF ATPase VmlR. Proc Natl Acad Sci U S A, 115, 8978–8983.

31. Su, W., Kumar, V., Ding, Y., Ero, R., Serra, A., Lee, B.S.T., Wong, A.S.W., Shi, J., Sze, S.K., Yang, L. et al. (2018) Ribosome protection by antibiotic resistance ATP-binding cassette protein. Proc Natl Acad Sci U S A, 115, 5157–5162.

32. Crowe-McAuliffe, C., Murina, V., Turnbull, K.J., Kasari, M., Mohamad, M., Polte, C., Takada, H., Vaitkevicius, K., Johansson, J., Ignatova, Z. et al. (2021) Structural basis of ABCF-mediated resistance to pleuromutilin, lincosamide, and streptogramin A antibiotics in Gram-positive pathogens. Nat Commun, 12, 3577.

33. Crowe-McAuliffe, C., Murina, V., Turnbull, K.J., Huch, S., Kasari, M., Takada, H., Nersisyan, L., Sundsfjord, A., Hegstad, K., Atkinson, G.C. et al. (2022) Structural basis for PoxtA-mediated resistance to phenicol and oxazolidinone antibiotics. Nat Commun, 13, 1860.

34. Cui, Z., Li, X., Shin, J., Gamper, H., Hou, Y.M., Sacchettini, J.C. and Zhang, J. (2022) Interplay between an ATP-binding cassette F protein and the ribosome from *Mycobacterium tuberculosis*. Nat Commun, 13, 432.

35. Romero, Z.J., Armstrong, T.J., Henrikus, S.S., Chen, S.H., Glass, D.J., Ferrazzoli, A.E., Wood, E.A., Chitteni-Pattu, S., van Oijen, A.M., Lovett, S.T., et al. (2020) Frequent template switching in postreplication gaps: suppression of deleterious consequences by the *Escherichia coli* Uup and RadD proteins. Nucleic Acids Res, 48, 212–230.

36. Romero, Z.J., Chen, S.H., Armstrong, T., Wood, E.A., van Oijen, A., Robinson, A. and Cox, M.M. (2020) Resolving Toxic DNA repair intermediates in every *E. coli* replication cycle: critical roles for RecG, Uup and RadD. Nucleic Acids Res, 48, 8445–8460.

37. Murat, D., Bance, P., Callebaut, I. and Dassa, E. (2006) ATP hydrolysis is essential for the function of the Uup ATP-binding cassette ATPase in precise excision of transposons. J Biol Chem, 281, 6850–6859.

38. Ousalem, F., Singh, S., Bailey, N.A., Wong, K.-H., Zhu, L., Neky, M.J., Sibindi, C., Fei, J., Gonzalez, R.L.J., Boël, G., et al. (2023) Comparative genetic, biochemical, and biophysical analyses of the four *E. coli* ABCF paralogs support distinct functions related to mRNA translation. bioRxiv, 2023.2006.2011.543863.

39. Mohamad, M., Nicholson, D., Saha, C.K., Hauryliuk, V., Edwards, T.A., Atkinson, G.C., Ranson, N.A. and O’Neill, A.J. (2022) Sal-type ABC-F proteins: intrinsic and common mediators of pleuromutilin resistance by target protection in staphylococci. Nucleic Acids Res, 50, 2128–2142.

40. Obana, N., Takada, H., Crowe-McAuliffe, C., Iwamoto, M., Egorov, A.A., Wu, K.J.Y., Chiba, S., Murina, V., Paternoga, H., Tresco, B.I.C. et al. (2023) Genome-encoded ABCF factors implicated in intrinsic antibiotic resistance in Gram-positive bacteria: VmlR2, Ard1 and CplR. Nucleic Acids Res, 51, 4536–4554.

41. Katoh, K. and Standley, D.M. (2013) MAFFT multiple sequence alignment software version 7: improvements in performance and usability. Mol Biol Evol, 30, 772–780.

42. Waterhouse, A.M., Procter, J.B., Martin, D.M., Clamp, M. and Barton, G.J. (2009) Jalview Version 2--a multiple sequence alignment editor and analysis workbench. Bioinformatics, 25, 1189-1191.

43. Capella-Gutierrez, S., Silla-Martinez, J.M. and Gabaldon, T. (2009) trimAl: a tool for automated alignment trimming in large-scale phylogenetic analyses. Bioinformatics, 25, 1972–1973.

44. Nguyen, L.T., Schmidt, H.A., von Haeseler, A. and Minh, B.Q. (2015) IQ-TREE: a fast and effective stochastic algorithm for estimating maximum-likelihood phylogenies. Mol Biol Evol, 32, 268–274.

45. Trifinopoulos, J., Nguyen, L.T., von Haeseler, A. and Minh, B.Q. (2016) W-IQ-TREE: a fast online phylogenetic tool for maximum likelihood analysis. Nucleic Acids Res, 44, W232–235.

46. Hoang, D.T., Chernomor, O., von Haeseler, A., Minh, B.Q. and Vinh, L.S. (2018) UFBoot2: Improving the Ultrafast Bootstrap Approximation. Mol Biol Evol, 35, 518–522.

47. Shimokawa-Chiba, N., Muller, C., Fujiwara, K., Beckert, B., Ito, K., Wilson, D.N. and Chiba, S. (2019) Release factor-dependent ribosome rescue by BrfA in the Gram-positive bacterium *Bacillus subtilis*. Nat Commun, 10, 5397.

48. Takada, H., Roghanian, M., Murina, V., Dzhygyr, I., Murayama, R., Akanuma, G., Atkinson, G.C., Garcia-Pino, A. and Hauryliuk, V. (2020) The C-Terminal RRM/ACT Domain Is Crucial for Fine-Tuning the Activation of ‘Long’ RelA-SpoT Homolog Enzymes by Ribosomal Complexes. Front Microbiol, 11, 277.

49. Chadani, Y., Niwa, T., Izumi, T., Sugata, N., Nagao, A., Suzuki, T., Chiba, S., Ito, K. and Taguchi, H. (2017) Intrinsic Ribosome Destabilization Underlies Translation and Provides an Organism with a Strategy of Environmental Sensing. Mol Cell, 68, 528–539 e525.

50. Britton, R.A., Eichenberger, P., Gonzalez-Pastor, J.E., Fawcett, P., Monson, R., Losick, R. and Grossman, A.D. (2002) Genome-wide analysis of the stationary-phase sigma factor (sigma-H) regulon of *Bacillus subtilis*. J Bacteriol, 184, 4881–4890.

51. Fujiwara, K., Katagi, Y., Ito, K. and Chiba, S. (2020) Proteome-wide Capture of Co-translational Protein Dynamics in Bacillus subtilis Using TnDR, a Transposable Protein-Dynamics Reporter. Cell Rep, 33, 108250.

52. Schneider, C.A., Rasband, W.S. and Eliceiri, K.W. (2012) NIH Image to ImageJ: 25 years of image analysis. Nat Methods, 9, 671–675.

53. Hummels, K.R. and Kearns, D.B. (2019) Suppressor mutations in ribosomal proteins and FliY restore Bacillus subtilis swarming motility in the absence of EF-P. PLoS Genet, 15, e1008179.

54. Pinheiro, B., Scheidler, C.M., Kielkowski, P., Schmid, M., Forne, I., Ye, S., Reiling, N., Takano, E., Imhof, A., Sieber, S.A. et al. (2020) Structure and Function of an Elongation Factor P Subfamily in Actinobacteria. Cell Rep, 30, 4332–4342 e4335.

55. Hummels, K.R., Witzky, A., Rajkovic, A., Tollerson, R., 2nd1, Jones, L.A., Ibba, M. and Kearns, D.B. (2017) Carbonyl reduction by YmfI in *Bacillus subtilis* prevents accumulation of an inhibitory EF-P modification state. Mol Microbiol, 106, 236–251.

56. Chandrasekaran, V., Juszkiewicz, S., Choi, J., Puglisi, J.D., Brown, A., Shao, S., Ramakrishnan, V. and Hegde, R.S. (2019) Mechanism of ribosome stalling during translation of a poly(A) tail. Nat Struct Mol Biol, 26, 1132–1140.

57. Koutmou, K.S., Schuller, A.P., Brunelle, J.L., Radhakrishnan, A., Djuranovic, S. and Green, R. (2015) Ribosomes slide on lysine-encoding homopolymeric A stretches. Elife, 4.

58. Kriachkov, V., Ormsby, A.R., Kusnadi, E.P., McWilliam, H.E.G., Mintern, J.D., Amarasinghe, S.L., Ritchie, M.E., Furic, L. and Hatters, D.M. (2023) Arginine-rich C9ORF72 ALS proteins stall ribosomes in a manner distinct from a canonical ribosome-associated quality control substrate. J Biol Chem, 299, 102774.

59. Lu, J. and Deutsch, C. (2008) Electrostatics in the ribosomal tunnel modulate chain elongation rates. J Mol Biol, 384, 73–86.

60. Shikha, S., Riley, C.G., Ziao, F., Nevette, A.B., Clara, G.A., Kam-Ho, W., Matthew, J.N., Chi, W., Korak Kumar, R., Ying-Chih, C., et al. (2023) Cryo-EM studies of the four *E. coli* paralogs establish ABCF proteins as master plumbers of the peptidyl-transferase center of the ribosome. bioRxiv, 2023.2006.2015.543498.

61. Yuhei, C., Eri, U., Kohei, Y., Miku, K. and Hideki, T. (2023) The ABCF proteins in *Escherichia coli* individually alleviate nascent peptide-induced noncanonical translations. bioRxiv, 2023.2010.2004.560807.

62. Ousalem, F., Saravuth, N., Thomas, O., Amin, O.-N., Marion, H., Laura, M. and Boël, G. (2023) Global regulation via modulation of ribosome pausing by EttA. bioRxiv, 2023.2010.2017.562674.

63. Hye-Rim, H., Cassidy, R.P., Letian, W. and Heather, A.F. (2023) Identification of factors that prevent ribosome stalling during early elongation. bioRxiv, 2023.2008.2004.552005.

64. Mohammad, F., Green, R. and Buskirk, A.R. (2019) A systematically-revised ribosome profiling method for bacteria reveals pauses at single-codon resolution. Elife, 8.

65. Huch, S., Nersisyan, L., Ropat, M., Barrett, D., Wu, M., Wang, J., Valeriano, V.D., Vardazaryan, N., Huerta-Cepas, J., Wei, W. et al. (2023) Atlas of mRNA translation and decay for bacteria. Nat Microbiol, 8, 1123–1136.

66. Katoh, T. and Suga, H. (2018) Ribosomal Incorporation of Consecutive beta-Amino Acids. J Am Chem Soc, 140, 12159–12167.

67. Daskalova, S.M., Dedkova, L.M., Maini, R., Talukder, P., Bai, X., Chowdhury, S.R., Zhang, C., Nangreave, R.C. and Hecht, S.M. (2023) Elongation Factor P Modulates the Incorporation of Structurally Diverse Noncanonical Amino Acids into *Escherichia coli* Dihydrofolate Reductase. J Am Chem Soc, 145, 23600–23608.

